# Uracil Repair - A Source of DNA Glycosylase Dependent Genome Instability

**DOI:** 10.1101/2023.03.07.530818

**Authors:** Claudia Krawczyk, Marc Bentele, Vitaly Latypov, Katrina Woolcock, Faiza Noreen, Olivier Fritsch, Primo Schär

## Abstract

Uracil DNA glycosylases (UDGs) excise uracil from DNA arising from dUMP misincorporation during replication or from cytosine deamination. Besides functioning in canonical uracil repair, UDGs cooperate with DNA base modifying enzymes to effect mutagenesis or DNA demethylation. Mammalian cells express four UDGs, the functional dissection of which represents a challenge. Here, we used *Schizosaccharomyces pombe* with only two UDGs, Ung1 and Thp1, as a simpler model to study functional interactions in uracil repair. We show that despite a predominance of Ung1 activity in cell extracts, both UDGs act redundantly against genomic uracil accumulation and mutations from cytosine deamination in cells. Notably, Thp1 but not Ung1-dependent repair is cytotoxic under genomic uracil stress induced by 5-fluorouracil exposure or AID expression. Also, Thp1-but not Ung1-mediated base excision is recombinogenic, accounting for more than 60% of spontaneous mitotic recombination events in a recombination assay. Hence, the qualitative outcome of uracil repair depends on the initiating UDG; while Ung1 shows expected features of a bona-fide DNA repair enzyme, Thp1-initiated repair appears slow and rather non-productive, suggesting a function beyond canonical DNA repair. Given the epigenetic role of TDGs, the mammalian orthologs of Thp1, we performed transcriptome analyses and identified a possible function of Thp1 in stabilizing gene expression.

## 1. Introduction

The intrinsic chemical instability of the DNA, as well as its error prone replication, constitutes significant threats to the stability of genomes. Uracil in DNA, for instance, arises by spontaneous deamination of cytosines or occasional dUMP misincorporation during DNA replication (1). While the former generates G•U mispairs that, unless corrected, will give rise to C→T transition mutations during DNA replication, misincorporated uracils are not mutagenic per se but can affect gene expression by interfering with transcription factor binding (2). Uracil and/or its derivatives are also generated by targeted enzymatic reactions for purposes such as genome editing during B cell activation (3-5), the restriction of retroviral replication (6, 7) and, possibly, active DNA demethylation (8, 9).

Uracil DNA glycosylases (UDGs) constitute a family of DNA repair enzymes that excise uracil and derivatives from DNA by hydrolyzing the *N*-glycosidic bond connecting the base with the sugar moiety of the DNA backbone. To restore the original DNA sequence, the resulting abasic site (AP-site) is then excised and replaced with a canonical nucleotide in a concerted enzymatic process known as DNA base excision repair (BER) (reviewed in (10)). Notably, UDGs have evolved in multiple flavors with different enzymatic properties and they show a species-specific co-existence, reflecting a biological need for variable tolerance, accumulation and processing of the non-canonical DNA base. Mammalian cells express four nuclear UDGs (11). UNG2 (uracil DNA glycosylase 2) is active on uracil and 5-fluorouracil (5-FU) and operates with an exceptionally high enzymatic turnover that facilitates efficient BER. It associates with the DNA replication apparatus where it provides the major activity for uracil excision following dUMP misincorporation (12-14). UNG2 also efficiently initiates BER of G•U mispairs at sites of cytosine deamination, but other UDGs, in particular Smug1 (single-strand-selective monofunctional uracil DNA glycosylase 1) appear to function redundantly in this context (15, 16). In addition to error-free repair, UNG2 is prominently involved in the mutagenic and recombinogenic processing of uracils generated by the activation-induced deaminase (AID) in B-cells, thereby contributing to somatic hypermutation (SHM) and class switch recombination (CSR). SMUG1 and other UDGs do not seem to engage in these processes but the evidence available does not strictly exclude the possibility (3, 17). The two remaining mammalian UDGs, TDG (thymine DNA glycosylase) and MBD4 (methyl CpG binding domain protein 4), are largely mismatch dependent and excise a broad spectrum of non-canonical bases (mis)paired with guanine (18, 19). While MBD4 was shown to counteract mutation at CpG dinucleotides (20), TDG appears to have only a minor role in mutation avoidance (21). Yet, the inverse cell-cycle-dependent regulation of TDG and UNG2 protein levels (22) indicates a primary function of TDG outside of S-phase and/or in non-dividing cells, in line with the increased mutation rates observed in stationary phase *E. coli* cells lacking the TDG ortholog Mug (23). The loss of TDG in mammalian cells increases their resistance to the chemotherapeutic drug 5-FU, which is a likely consequence of its rate-limiting dissociation from AP-sites and, hence, an accumulation of this labile repair intermediate under UNG2 saturating conditions (24). Strikingly, TDG is the only DNA glycosylase essential for embryonic development. It is required for proper gene expression and setting of epigenetic marks during embryogenesis (21, 25) and acts as a co-regulator of nuclear receptors (26-28). TDG, but also MDB4, have been implicated in the active demethylation of 5-methylcytosine (5mC) (reviewed in (11)), which involves excision of oxidation and/or deamination products generated by the ten eleven translocation (TET) (29-31) or the AID/APOBEC proteins (25, 32, 33). Hence, UDGs operate in diverse and complex processes in mammalian cells, making it difficult to dissect their specific biological roles and functional interactions.

The fission yeast *Schizosaccharomyces pombe* encodes only two UDGs, the UNG2 ortholog Ung1 and the TDG ortholog Thp1. Ung1 was shown to be a highly active UDG capable of excising uracil from DNA *in vitro* (34). Although disruption of *ung1*^*+*^ in *S. pombe* cells did not reveal any obvious phenotypes, overexpression caused cell toxicity accompanied by an accumulation of AP-sites and increased mutation frequencies (34). Like its human counterpart, Ung1 interacts with proliferating cell nuclear antigen (PCNA) (35) and is therefore likely to act in the context of DNA replication. Thp1 was shown to be active on a broad range of substrates (36). Yet, unlike its mammalian orthologs and in line with the lack of DNA methylation in fission yeast (37), it is incapable of excising the 5mC derivatives thymine and 5hmU opposite guanine (36). Similar to its mammalian homolog, however, a physical interaction with the gene activator protein Cbf11, identified in a proteomic screen, implicates a potential role of Thp1 in gene regulation (38, 39).

We used *S. pombe* to address the functional and mechanistic interactions of UDGs in a system of reduced complexity. Investigating the characteristics of Ung1- and Thp1-dependent DNA repair both genetically and biochemically, we found that both UDGs act redundantly in the elimination of genomic uracil and in avoiding mutation by cytosine deamination. This is remarkable in the light of Ung1 constituting the only detectable uracil excision activity in cell-free extracts. Challenging the cells with high levels of genomic uracil either by 5-FU treatment or by overexpression of AID, however, revealed that the two UDGs operate with different modes of action, generating different repair outcomes; repair by Thp1 caused cell death under these conditions while repair by Ung1 was generally protective. The cytotoxic effects likely reflect the Thp1-dependent formation of long-lived AP-sites in DNA. Consistently, Thp1-induced repair was also recombinogenic, accounting for more than 60% of spontaneous and X-ray induced mitotic recombination. Given this unproductive role in DNA repair and given the epigenetic function of its mammalian orthologs, we explored a potential role of Thp1 in the regulation of gene expression. Genome-wide analyses revealed a slight but overall suppression of gene expression in Thp1-deficient cells, which was accompanied with a higher variation of transcript levels across replicas. These findings indicate a potential function of Thp1 in maintaining transcriptionally active chromatin.

## 2. Material and methods

### 2.1. Strains, growth conditions, plasmids, oligonucleotides

All strains used in this study derive from the 972 *h*^*-*^ wild-type strain (Supplementary Table 1). Gene disruption and tagging were achieved by using PCR generated fragments providing short homology for recombination as described (40). Complete *thp1*^*+*^ or *ung1*^*+*^ open reading frames were replaced in strain PRS000d. Enhanced green fluorescent (EGFP) tagging of *thp1*^*+*^ was obtained by first inserting a *ura4*^*+*^ gene 3’ of the *thp1*^*+*^ open reading frame in the PRS000d strain that was subsequently replaced by the EGFP coding sequence.

Standard media and growth conditions were described before (41). EMM-Can-G plates contained 3.75 g/l of glutamate as the nitrogen source and 75 μg/ml of L-canavanine sulphate (Sigma). For inducible *nmt1* promoter driven overexpression (42), cells were initially grown under repressive conditions (EMM + 5 μg/ml thiamine) to 1×10^7^ cells/ml, washed twice in water, diluted for induction with EMM lacking thiamine to a density of 5×10^5^ cells/ml and grown at 30°C, 16 h.

pPRS271 and pREP1-AID for Thp1 and AID overexpression, respectively, were constructed by sub-cloning the *thp1*^*+*^ and human *AID* open reading frames as NdeI-SalI PCR fragments into the matching sites of pREP1 (ARS1/*LEU2* based episomal *S. pombe* vector for inducible *nmt1* promoter-driven gene expression). Oligonucleotide sequences are available on request.

### 2.2. Cell sensitivity tests

To test cellular 5-FU sensitivity, cells were grown to 5×10^6^ cells/ml and washed in water. Ten fold serial dilutions were spotted onto MMA plates supplemented with the indicated concentration of 5-FU and incubated for 6-12 days at 26°C. To test sensitivity to AID overexpression, cells transformed with pREP1-AID were grown under repressive conditions (EMM + 5 μg/ml thiamine) to a density of 5×10^6^ cells/ml, harvested and washed in water. Serial dilutions of cells in water were spotted onto EMM (inducible conditions) and incubated for 7 days at 30^°^C.

### 2.3. Cell-free extracts, Western blotting and base release assay

Cells were harvested at 1×10^7^ cells/ml and sequentially washed in water and twice in 10 ml of lysis buffer (50 mM Tris-HCl pH 8.0, 500 mM NaCl, 1 mM EDTA, 20% glycerol, 0.1% Tween-20, 10 mM β-mercaptoethanol, 1 mM PMSF, 1x complete™ protease inhibitors (Roche)). Cells were resuspended in ice-cold lysis buffer and disrupted by adding glass beads and vigorous shaking for ten times 30 s in a Mini-Beadbeater (Biospec Products). After centrifugation (20800 g, 20 min, 4°C), protein concentrations of the supernatant were determined using a Bradford assay. For Western blotting, 50 μg of protein were separated on 10% SDS-polyacrylamide gels, transferred to a nitrocellulose membrane (Protran®, Schleicher & Schuell) and incubated with an affinity purified rabbit polyclonal anti-Thp1 antiserum or a mouse anti-AID antiserum according to standard protocols. The rabbit polyclonal antiserum was raised against purified recombinant Thp1 protein and subsequently affinity purified as Thp1p coated affinity beads (Affi-Gel 10 beads, Bio-Rad) according to standard procedures. This Thp1 antibody was diluted 1:500 in TBS-T containing 5% dry milk as blocking reagent.

For base release assays, a 6His-tagged Thp1 was expressed in *E. coli* and purified as described (36). Substrate preparation and nicking assay were performed as previously described ((43), see Supplementary Methods for details).

### 2.4. Mutation and intra-chromosomal recombination rates, mutation spectra

Spontaneous recombination rates were determined at the *ade6-M387* allele (44). For fluctuation tests, at least 24 colonies freshly grown on non-selective medium (YEA supplemented with adenine and uracil) were used to inoculate 4 ml of fresh non-selective medium. After 24 h of incubation at 30°C, cells were harvested, washed with water and resuspended in 3 ml of water. 1 ml of the cell suspension was plated onto five MMA plates supplemented with uracil. Adenine prototrophic colonies (revertants) were scored after 9-11 days at 30°C. For mutation rates upon overexpression of Thp1, three experiments with four cultures each were performed. To measure rates of spontaneous forward mutation conferring canavanine resistance (45), 30 independent cultures from 3 experiments were used for fluctuation analysis. Cells from single colonies, freshly grown on non-selective YEA plates, were grown in 5 ml YEL for 28 h at 30°C, harvested, washed in water, resuspended in 1.2 ml of water and plated onto 4 selective EMM-Can-G plates (0.2 ml/plate). Colonies were scored after 14-16 days at 30°C. For 5-FU-induced mutation rates, at least 3 independent experiments were performed in which one culture per strain was split in two halves and grown to midlog phase in YEL. One half culture was supplemented with a final concentration of 10 mg/l 5-FU and both were incubated for 76 h at 30°C before being processed and analyzed as above. Inducibility of canavanine resistance was calculated as follows: [(induced mutations of mutant) x (spontaneous mutations of wild-type)] / [(spontaneous mutations of mutant) x (induced mutations of wild-type)].

To measure intra-chromosomal recombination, heteroallelic duplication of *ade6*^*+*^ mutant alleles (*ade6-L469, ade6-M375*, (46)) were crossed into our strain backgrounds. One freshly grown red (adenine auxotrophic) colony was isolated from YEA plates and re-suspended in water. 100 cells were plated onto two YEA plates and incubated for 4 days at 30°C. At least 24 randomly chosen colonies (excluding white colonies) were re-suspended in a final volume of 0.7 ml. For IR induced recombination analyses, cells suspended in H_2_O were X-ray exposed (100Gy, 100kv, 0.2mm Al) before being plated onto YEA. Two times 0.2 ml were spread onto EMM plates supplemented with uracil. Adenine prototrophs were scored after 3 days at 30°C.

The number of viable cells was determined on non-selective YEA plates after 3 days at 30°C. Spontaneous reversion rates were determined by the method of the median (47). The 95% confidence interval for the median was calculated according to Nair (48). To determine the mutation spectra among the spontaneous *ade6-M387* revertants, one randomly chosen adenine prototrophic clone per culture was used for sequence analysis. Transversion and transition rates were calculated using the following formula: (transversions or transitions) x (reversion-rate) / (number of tested clones).

### 2.5. Fluorescence microscopy

Cells were arrested by glucose starvation for 16 h in EMM containing 0.5% glucose. 500 μl of cells were harvested, washed in water, resuspended in 1 M sorbitol and stained with Hoechst 33342 dye (1 μg/ml, Sigma). Images of EGFP and Hoechst 33342 were analyzed by standard epi-fluorescence microscopy.

### 2.6. DNA isolation in agarose plugs and PFGE

Cells were grown in YEL to a density of 1×10^7^ cells/ml at 30°C. For 5-FU treatment, cells were grown in YEL to 5×10^6^ cells/ml before 5-FU was added to a final concentration of 10 mg/l for 48 h at 30°C. DNA isolation in agarose plugs and PFGE were done as in (49) with some modifications (see Supplementary Methods).

### 2.7. Genome-wide gene expression profiling

WT (975 *h*^*+*^) and *thp1Δ* (PRS555) strains were crossed to isolate 9 WT and 9 *thp1Δ* spore clones with an *h*^*+*^ mating type. RNA isolation was done with acidic (pH 4.3) phenol as described previously (50). RNAs from three independent clones were pooled and three such pools for each genotype were analyzed on Affymetrix GeneChip^®^ *S. pombe* Tiling 1.0FR Arrays as previously described (51). Linear regression analyses and the Pearson correlation coefficient calculation were performed using GraphPad Prism (version 6.0 for Mac OS X, GraphPad Software). Principal component analysis was performed using Cluster 3.0 and the results visualized in TOPCAT (http://www.star.bristol.ac.uk/~mbt/topcat/).

## 3. Results

### 3.1. Ung1 constitutes the major uracil excision activity in S. pombe extracts

To assess the relative contributions of Ung1 and Thp1 to the uracil processing activity present in *S. pombe* cells, we first measured efficiencies of uracil removal from A•U base pairs and G•U mismatches in cell-free extracts (43). As substrates, we used synthetic, 5’-fluorescein labeled 60-mer DNA duplexes containing a single uracil paired with either an adenine or a guanine (Fig. 1A). Uracil excision from these substrates followed by AP-site cleavage generates a 23-mer product fragment. As a reference, we included recombinant Thp1, which we showed previously to excise uracil opposite from adenine and guanine (36). Extracts of wild-type cells showed high uracil excision activity on both substrates. No activity was discernible with extracts from *thp1*Δ*ung1*Δ double mutant cells, indicating that Ung1 and Thp1 account for all detectable UDG activity in *S. pombe*. To distinguish Ung1 from Thp1 activity in wild-type extracts, we inhibited Ung1 by adding Ugi peptide, a specific and potent inhibitor of the Ung but not the Mug family of UDGs (36, 52). Addition of Ugi eliminated all detectable uracil processing, both on A•U and G•U substrates, despite the presence of appreciable levels Thp1 in exponentially growing cells (Fig. 1A and 1B). Based on a detection limit of 5% of total substrate used in this assay, we conclude that Ung1 accounts for more than 95% of uracil excision activity in wild-type cell extracts. Under fully Ung1 inhibited conditions, only a strong ectopic expression of *thp1*^*+*^ from the inducible *nmt1* promoter (Fig. 1B) yielded a robust uracil excision activity on both A•U and G•U substrates. Hence, Thp1 has the ability to excise uracil from DNA but, compared to Ung1, it contributes only marginally, if anything, to the uracil processing in extracts.

**Fig. 1.**
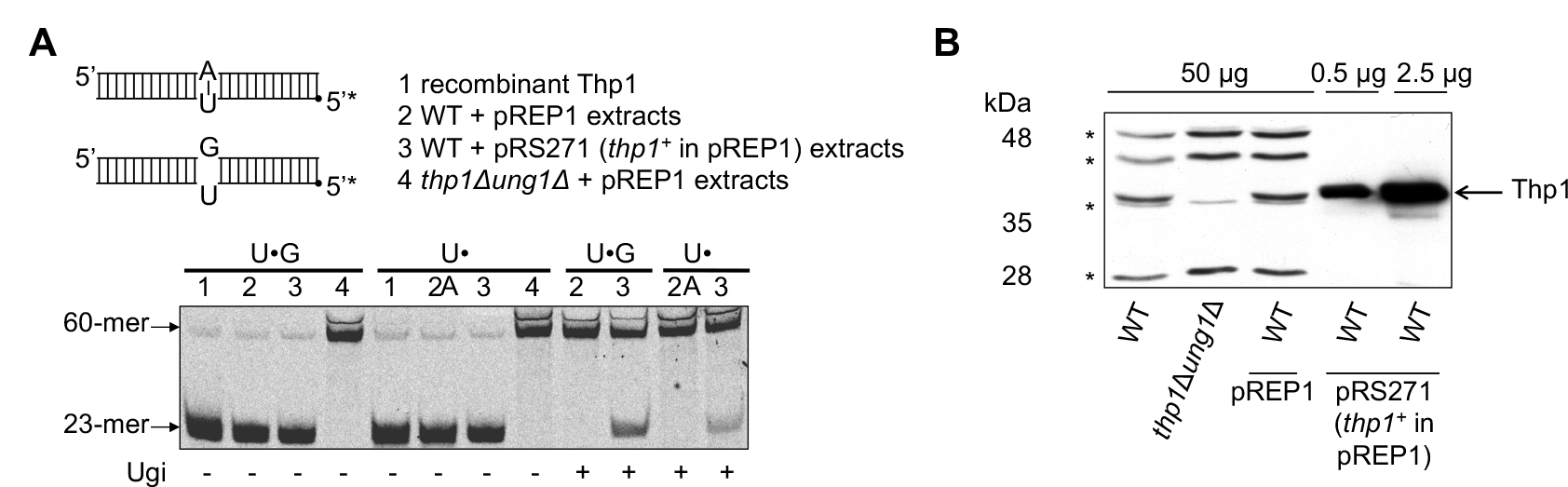
Ung1 constitutes the main uracil excision activity in cell extracts. **(A)** DNA nicking assay with a 60-mer 5’ fluorescein labeled (*) 60-mer double-stranded DNA substrate containing an A•U base-pair or a G•U mismatch. Uracil excision results in a labeled 23-mer upon AP-site hydrolysis. 2 pmol of substrate were incubated with 2 pmol of recombinant Thp1 or 7 μg of cell-free extracts of the strains indicated; Uracil-DNA glycosylase inhibitor (Ugi, 1U) was added as indicated. For Thp1 overexpression, cells transformed with pPRS271 or with the vector pREP1 were induced for 16 h. **(B)** Crude protein extracts of the strains indicated and induced as in (A) were prepared and analyzed by immunoblotting with an affinity purified rabbit polyclonal anti-Thp1 antiserum. Arrow, Thp1; *, unspecific bands (*).

### 3.2. Ung1 and Thp1 both remove uracil from genomic DNA

While the nuclear localization of Ung1 was demonstrated earlier (34), we determined the subcellular localization of Thp1 by expressing a C-terminally tagged Thp1-EGFP protein from the endogenous *thp1+* locus. This fusion protein produced a distinct nuclear EGFP signal (Fig. 2A), consistent with the presence of two predicted nuclear localization signals (NLSs) (53) and public localization data (http://www.riken.jp/SPD/Img_page/14_iP/14A10_Loc.html).

**Fig. 2.**
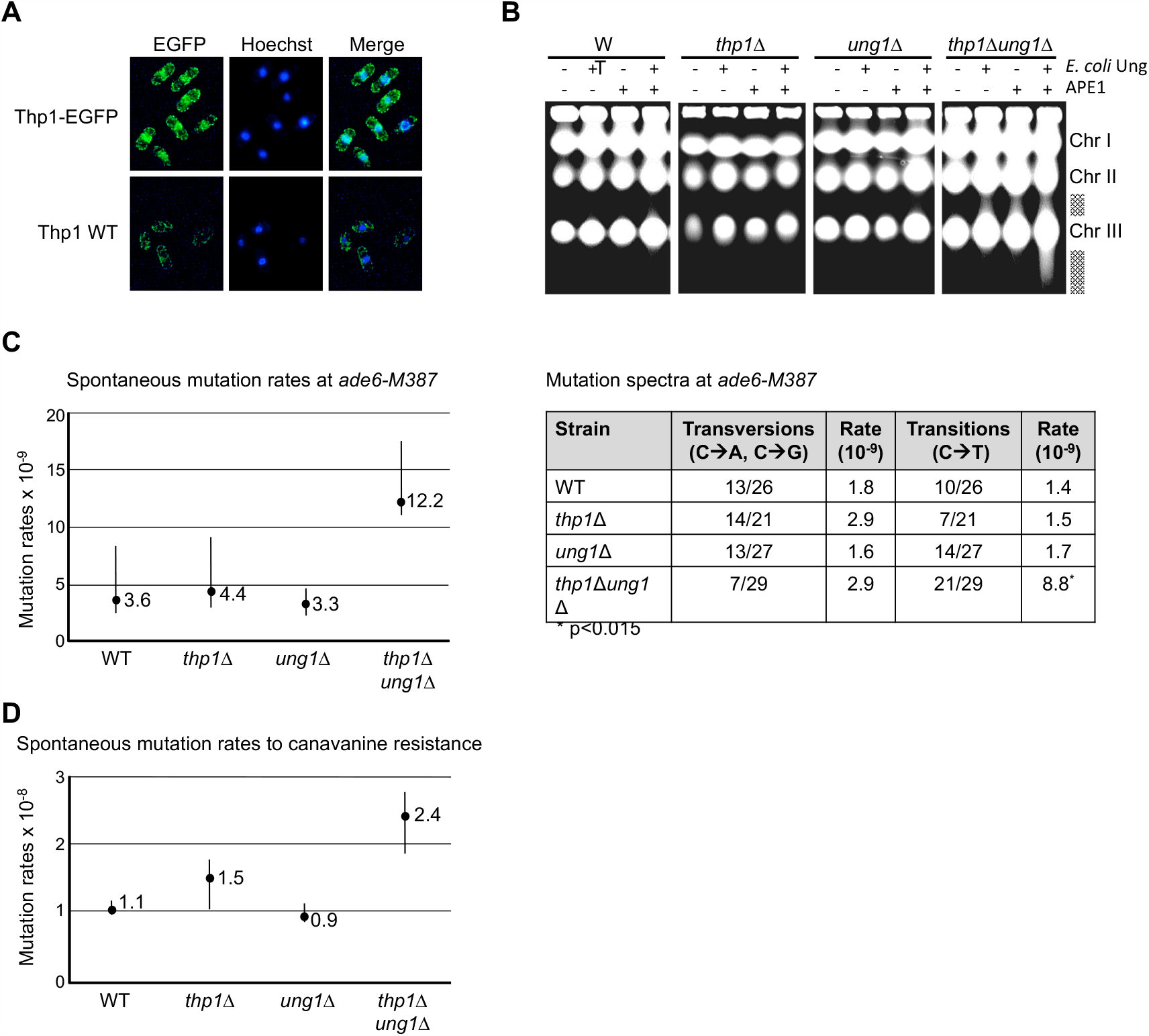
Thp1 and Ung1 cooperate in anti-mutagenic uracil repair. **(A)** Subcellular localization of Thp1 in wild-type cells. Fluorescence microscopy of cells expressing a C-terminally tagged Thp1-EGFP fusion protein from the endogenous *thp1* locus or the untagged control. **(B)** Uracil accumulation in *thp1*Δ *ung1*Δ strains. Shown is chromosomal DNA prepared in agarose plugs, digested with *E. coli* Ung and/or human AP-endonuclease (APE1) as indicated, separated by PFGE and stained with ethidium bromide. Intact *S. pombe* chromosomes (Chr) I, II and III and fragmented DNA (shaded bar) are indicated. **(C)** Spontaneous reversion rates and mutation spectra at *ade6-M387*. Any base substitution at *ade6-M387* restores adenine prototrophy. Shown are mutation rates with 95% confidential intervals as calculated from prototrophs arising in at least 24 independent cultures out of 3 experiments. Corresponding mutation spectra, determined by DNA sequencing of one random clone of each culture, are shown in the graph, statistical significance (*) was assessed by the Fisher’s exact test. **(D)** Spontaneous forward mutation rates to L-canavanine resistance. Clones with mutations at the arginine permease gene *can1*^*+*^ were scored after 14 to 16 days on EMM plates containing L-canavanine. Mutation rates were determined using at least 30 independent cultures out of 3 experiments. Shown are the median rates with 95% confidential intervals.

Given that both UDGs are expressed and localize to the nucleus in *S. pombe* cells, we next addressed their individual roles in uracil removal. We first assessed the uracil content in genomic DNA isolated from exponentially growing wild-type, Ung1-, Thp1- and doubly-deficient cells. To this end, we extracted intact DNA in agarose plugs, digested it with *E. coli* Ung uracil-DNA glycosylase and/or human AP-endonuclease APE1 to excise potential uracils and incise the resulting AP-sites, and finally analyzed the products by pulse field electrophoresis (PFGE). This treatment is expected to generate single-stranded DNA breaks at uracil residues, which, if closely spaced on opposite strands, will give rise to DNA fragmentation that can be visualized by PFGE (Fig. 2B). Indeed, substantial smearing of the chromosomal bands was apparent in the DNA derived from *thp1*Δ*ung1*Δ double mutant cells digested with both Ung and APE1. DNA from either single mutant or wild-type cells at the same conditions remained intact during digestions. From this result we conclude that uracil arises in DNA of vegetatively growing *S. pombe* cells and, as this is detectable mainly in double mutant cells, that both Ung1 and Thp1 act redundantly in uracil excision repair.

### 3.3. Lack of uracil repair generates a moderate mutator phenotype

We wondered whether the accumulation of uracil in the absence of Ung1 and Thp1 coincides with increased mutation as expected if a sizable fraction of these uracils resulted from cytosine deamination. We assessed mutation rates and spectra in wild-type, *ung1*Δ and *thp1*Δ single mutant and *thp1*Δ*ung1*Δ double mutant cells. In a first assay, we scored mutations reversing the adenine auxotrophy of the *ade6-M387* allele, a G→C transversion that renders the *ade6* encoded protein non-functional but can be reverse by any of the three possible base substitution (C→G, C→A, C→T) (44). Fluctuation analyses revealed an increase of the spontaneous reversion rate by 3.4-fold in *thp1*Δ*ung1*Δ cells as compared to the wild-type, with non-overlapping 95% confidence intervals indicating statistical significance of the rate differences (Fig. 2C, left panel). Importantly, disruption of only *ung1*^*+*^ or *thp1*^*+*^ did not increase mutation rates, indicating that the two UDGs can fully compensate for each other in the defense against cytosine mutagenesis. Analysis of the mutation spectra by DNA sequencing of one randomly chosen adenine prototrophic clone per culture (to avoid effects of clonal amplification) unveiled an underlying 6.3-fold increase specifically in the C→T transition rate in *thp1*Δ*ung1*Δ double mutants relative to the wild-type (p=0.015, Fisher’s exact test; Fig. 2C, right panel). Neither of the single mutant showed a significant change in mutation spectrum. This result suggests that Thp1 and Ung1 operate synergistically in the uracil removal from pre-mutagenic G•U mispairs in cells. To corroborate the synergistic interaction of Ung1 and Thp1 in mutation avoidance, we performed a forward mutation assay, scoring for canavanine resistance (45). Again, *thp1*Δ*ung1*Δ double mutant cells but neither of the single mutants showed a moderate mutator phenotype when compared to wild-type cells (Fig. 2D). The observed 2.3-fold increase in forward mutation rate was statistically significant by the criterion of non-overlapping 95% confidence intervals. From these results, we conclude that Thp1 and Ung1 have redundant functions in the repair of cytosine deamination damage in cells.

### 3.4. Ung1 and Thp1 operate differently on 5-FU and AID induced DNA lesions

To further explore potential non-redundant functions of the two UDGs, we assessed the effect of *thp1*^*+*^ and *ung1*^*+*^ deletion under conditions of uracil stress, such as induced by exposure of cells to the anticancer drug 5-FU. 5-FU exerts its cytotoxic effects by inhibiting the thymidylate synthase that converts dUMP to dTMP, thereby leading to an imbalanced nucleotide pool and, consequently, increased dUMP incorporation into DNA. Moreover, 5-FU and its metabolites can be directly incorporated into both DNA and RNA (24, 54-56). 5-FU in DNA is a substrate for UDGs and will thus trigger repair events, whereas the consequences of 5-FU in RNA are not entirely clear. To ascertain 5-FU dependent uracil incorporation into *S. pombe* DNA, we digested genomic DNA of 5-FU treated wild-type cells with uracil DNA glycosylase and/or AP-endonuclease and examined the fragmentation of chromosomal DNA by PFGE. The pattern of DNA fragmentation observed indicated an accumulation of AP-sites (APE1 digest) in DNA of 5-FU treated but not untreated wild-type cells (Fig. 2B and 3A). No further increase in fragmentation was apparent in doubly-digested DNA (Ung and APE1), suggesting that AP-site processing rather than uracil excision is rate limiting in under these conditions. Consistently, we noticed a 5-FU dependent growth inhibition in wild-type cells (Fig. 3B). Notably, the 5-FU response of the two UDG single mutants was diametrically opposite. While Thp1-deficient cells displayed a striking hyper-resistance towards 5-FU, Ung1-deficient cells were hyper-sensitive at the applied doses. Consistent with this 5-FU sensitizing effect of Thp1, we observed that Thp1 overexpression from the inducible *nmt1* promoter strongly increases 5-FU sensitivity in wild-type and *ung1*Δ*thp1*Δ double mutant cells (Fig. 3C). In the double mutant configuration, the 5-FU hyper-sensitivity caused by the loss of Ung1 is largely suppressed by the hyper-resistance caused by Thp1 loss. Thus, 5-FU sensitivity of fission yeast is mediated by Thp1, while Ung1 promotes cell survival.

**Fig. 3.**
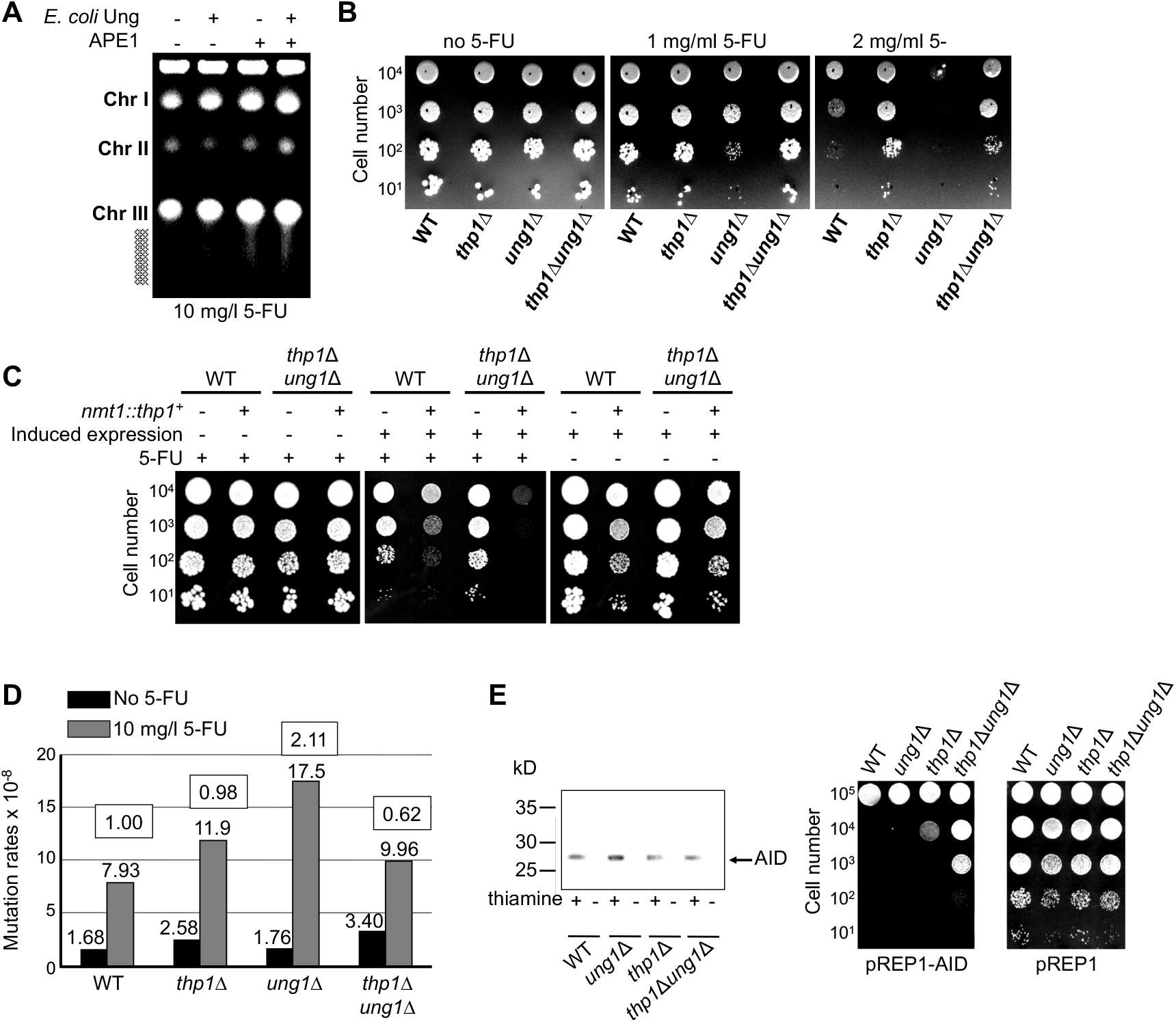
Thp1 sensitizes cells to 5-FU and ectopic expression of AID. **(A)** 5-FU exposure causes AP-site accumulation in wild-type cells. Chromosomal DNA from wild-type cells grown for 48 h in YEL supplemented with 10 mg/l 5-FU was digested with *E. coli* Ung and/or APE1 as indicated, separated by PFGE and stained with ethidium bromide. Fragmented DNA forms a smear (shaded bar) emanating from intact chromosome bands (Chr). **(B)** Cellular sensitivity to 5-FU. Serial dilutions of cells were spotted on MMA plates containing 5-FU as indicated and incubated for 6 to 12 days at 26°C. **(C)** Effect of Thp1 overexpression on 5-FU sensitivity. Serial dilutions of WT and *thp1Δ ung1Δ* double mutant cells transformed with control (pREP1) or Thp1 expression plasmids (PRS271) were spotted on EMM with (repressed expression) or without 5 μg/ml thiamine (induced expression) and incubated for 8 to 12 days at 26°C. **(D)** 5-FU induced forward mutations to canavanine resistance. Cultures were divided in halves and incubated for 76 h at 30°C in the absence or presence of 10 mg/l 5-FU. Spontaneous and induced mutation rates were determined from 3 to 4 cultures per strain. WT-normalized inducibility of mutations is indicated in white boxes. **(E)** Expression of human AID in *S. pombe* cells. Human AID was expressed from the pREP1 carrying the inducible *nmt1* promoter (pREP1-AID). Cells were grown under inducing (+) or repressing (-) conditions for 16 h. Crude protein extracts (50 μg) were separated on SDS-PAGE and subjected to Western blot analysis using mouse anti-AID antiserum; AID is indicated. **(F)** Cytotoxicity of 5-FU treatment. Serial dilutions of cells transformed with pREP1 or pREP1-AID were spotted onto EMM medium (inducible condition) and incubated for 7 days at 30°C.

We further investigated the effect of Thp1 expression on 5-FU induced mutations, assessing the rates of spontaneous and 5-FU induced mutations towards canavanine resistance (Fig. 3D). Ung1-deficient cells displayed the highest mutation inducibility upon 5-FU exposure, indicating that Ung1 controlled DNA repair reduces the mutagenic effect of 5-FU. Notably, the hyper-inducibility of mutations in Ung1-deficient cells depended on functional Thp1 as it was fully suppressed in the *ung1*Δ*thp1*Δ double mutant. From these results, we conclude that Ung1- and Thp1-mediated uracil and 5-FU repair operate through distinct pathways, the former increasing viability and suppressing mutations, the latter being cytotoxic and mutagenic.

To rationalize the differential response of Ung1- and Thp1-deficient cells to 5-FU, we reasoned that, due to its high affinity for AP-sites (36), Thp1 will generate AP-sites that are poorly accessible for further processing by the repair system. Hence, in Ung1 saturating conditions (5-FU exposure), Thp1-generated AP-sites would accumulate and trigger cell death and mutations. To test this hypothesis, we challenged cells by expression of the human activation-induced cytidine deaminase (AID), which will artificially generate G•U mismatches across the genome (Fig. 3E, left panel). Ectopic AID expression, driven by the inducible *nmt1*-promoter, induced severe cell death in wild-type, Ung1, Thp1 and doubly-deficient cells (Fig. 3E, right panel). Yet, the *ung1*Δ*thp1*Δ double mutants showed a more than 100-fold higher resistance to AID-expression than wild-type cells. This shows that uracil excision and, hence, the widespread generation of AP-sites, is a major cause of AID triggered cell death. Notably, AID toxicity was less pronounced in Thp1-than in Ung1-deficient cells, indicating that repair of G•U mismatches by Thp1 is more detrimental than repair by Ung1. We thus conclude that the repair of excessive uracil in DNA by UDGs generates toxic intermediates that are less efficiently processed when Thp1 rather than Ung1 is the initiating glycosylase.

### 3.5. Thp1-initiated BER is recombinogenic

We reasoned that the slow dissociation of Thp1 from AP-sites is responsible for its toxic effects during 5-FU exposure and AID expression. The generation of long-lived labile AP-sites is likely to be accompanied by occasional DNA strand-breakage, which would then trigger recombination events. To study a potential relationship between Thp1 initiated BER and recombination, we crossed a heteroallelic duplication of *ade6* mutant alleles (ade6-L469, ade6-M375, (46)) into our UDG-proficient and -deficient strain backgrounds. This system allows homologous recombination events. restoring an intact *ade6*^*+*^ gene. to be scored by selection for adenine prototrophy (*ade*^*+*^, Fig. 4A). We first determined spontaneous mitotic recombination rates in vegetatively growing cells (Figure 4B) and found that both wild-type and *ung1*Δ cells recombine the *ade6* alleles with similar rates (overlapping 95% confidence intervals). Thp1-deficient cells, however, displayed significantly reduced mitotic recombination; the rates with *thp1*Δ single and *ung1*Δ*thp1*Δ double mutant cells were more than 60% lower than those of wild-type cells. Therefore, Thp1-but not Ung1-dependent base excision is responsible for a significant part of spontaneously occurring mitotic recombination at the *ade6* locus.

**Fig. 4.**
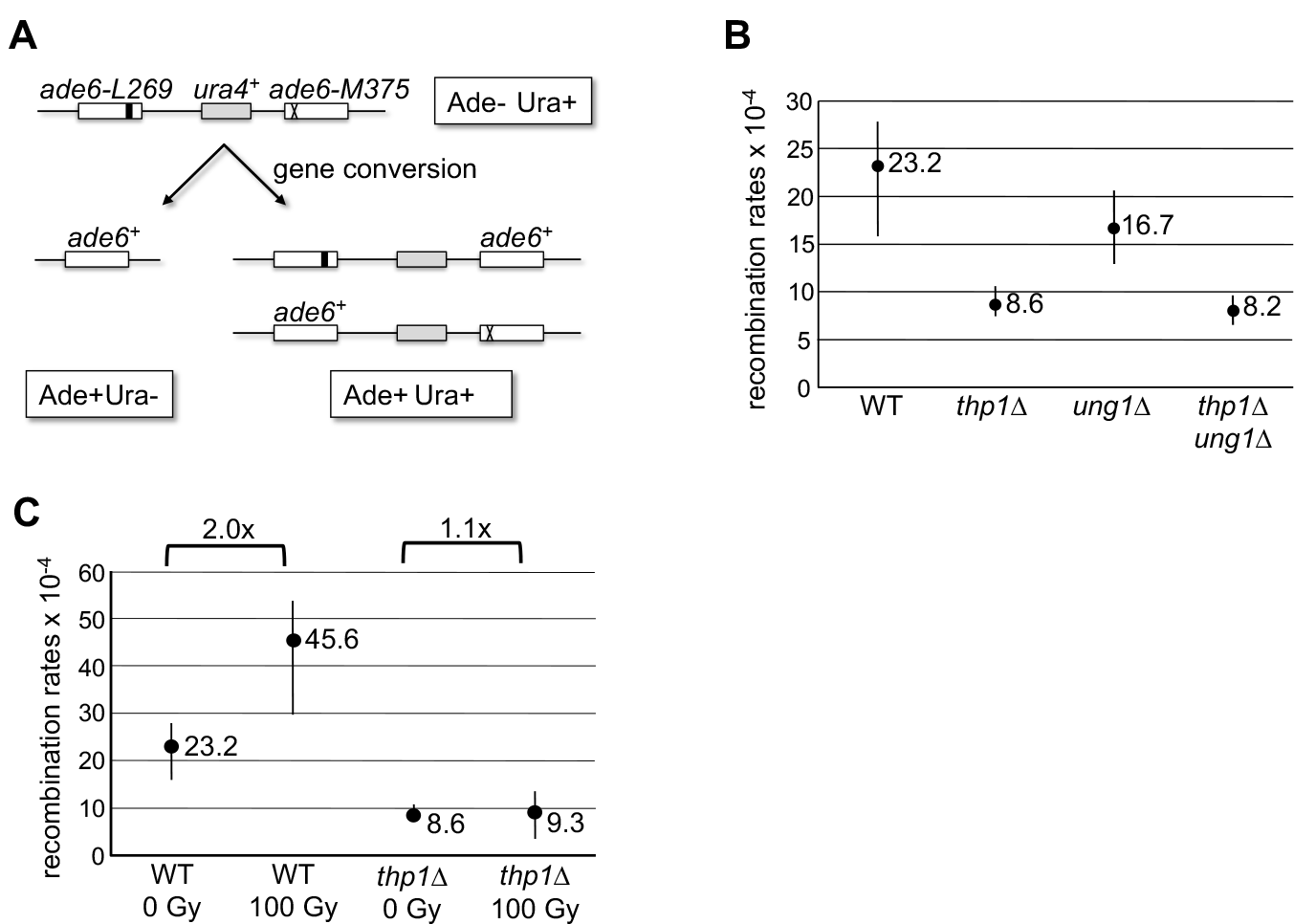
Thp1 triggers spontaneous and X-ray-induced recombination. **(A)** Scheme of the ectopic homologous recombination recombination assay, consisting of direct repeats of the *ade6-L469* and *ade6-M375* alleles, separated by a functional *ura4*^*+*^ gene. The assay scores for gene conversion events conferring adenine prototrophy. **(B)** Rates of spontaneous mitotic recombination. Recombination was scored by adenine prototrophy and recombination rates were calculated using at least 24 cultures per strain. Shown are median rated with 95% confidential intervals. **(C)** X-ray induced mitotic recombination. Cells were exposed to a non-lethal X-ray dose (100 Gy) before recombination recombination rates were determined as in (B). Shown are median rates with 95% confidential intervals.

To corroborate the recombinogenic action of Thp1, we determined its contribution to recombination under damage inducing conditions. For this purpose, we exposed wild-type and Thp1-deficient cells to a non-lethal dose of X-rays (100 Gy, >95% cell survival), which we expected to generate ROS-mediated Thp1 relevant DNA base damage, such as oxidized pyrimidines (Fig. 4C). Under these conditions, we measured a significant 2-fold increase in recombination rates in wild-type cells while no increase was apparent in Thp1-deficient cells. Hence, both, spontaneous and X-ray induced mitotic recombination show a clear Thp1 dependency. We thus conclude that repair of base-damage by Thp1 is recombinogenic and can result in gross genomic instability.

### 3.6. Effect of Thp1 on global gene expression

Considering its suboptimal performance in DNA repair, we wondered whether Thp1, similar to its mammalian ortholog TDG, could have evolved for purposes other than the conventional repair of DNA base damage, i.e. to control gene expression (21, 25). To address this possibility, we determined the genome-wide transcript levels of wild-type and *thp1*Δ strains with an *h*^*+*^ mating type (57). We isolated nine spores of each genotype from a cross between wild-type and *thp1*Δ cells and extracted RNA from cultures expanded from these single spores. We then pooled RNAs of three independent cultures for each genotype and used three of these pools for gene expression analysis on an Affymetrix GeneChip^®^ *S. pombe* Tiling 1.0FR Array. Comparing the mean log_2_ gene expression values of wild-type and Thp1-deficient cells revealed only minor differences in mRNA levels (Fig. 5A) as well as in separately analyzed transcript levels of 5’ and 3’ untranslated regions (5UTR, 3UTR, respectively) and long non-coding RNAs (lncRNAs, Supplementary Fig. 1A). Hence, under vegetative growth conditions, loss of Thp1 does not cause a specific deregulation of gene expression. However, Thp1-deficient cells displayed a general suppression of transcript levels when compared to wild-type cells (log_2_ fold change *thp1Δ*/wild-type, log_2_FC) affecting mRNAs (Fig. 5B), 5UTRs, 3UTRs and lncRNAs (Supplementary Fig. 1B); 96% of all differentially expressed mRNAs (log_2_FC *thp1Δ*/wild-type >0.3 or <-0.3, n= 471) were less expressed in Thp1-deficient cells (Fig. 5C). The high proportion of genes showing reduced transcription upon *thp1*^*+*^ deletion was not an effect of the applied threshold, it was also observed when all mRNAs were analyzed (Supplementary Fig. 1C). Although the individual expression differences were small and not statistically significant, together they indicate a consistent trend towards lower gene expression across most of the transcriptome in *thp1*Δ cells.

**Fig. 5.**
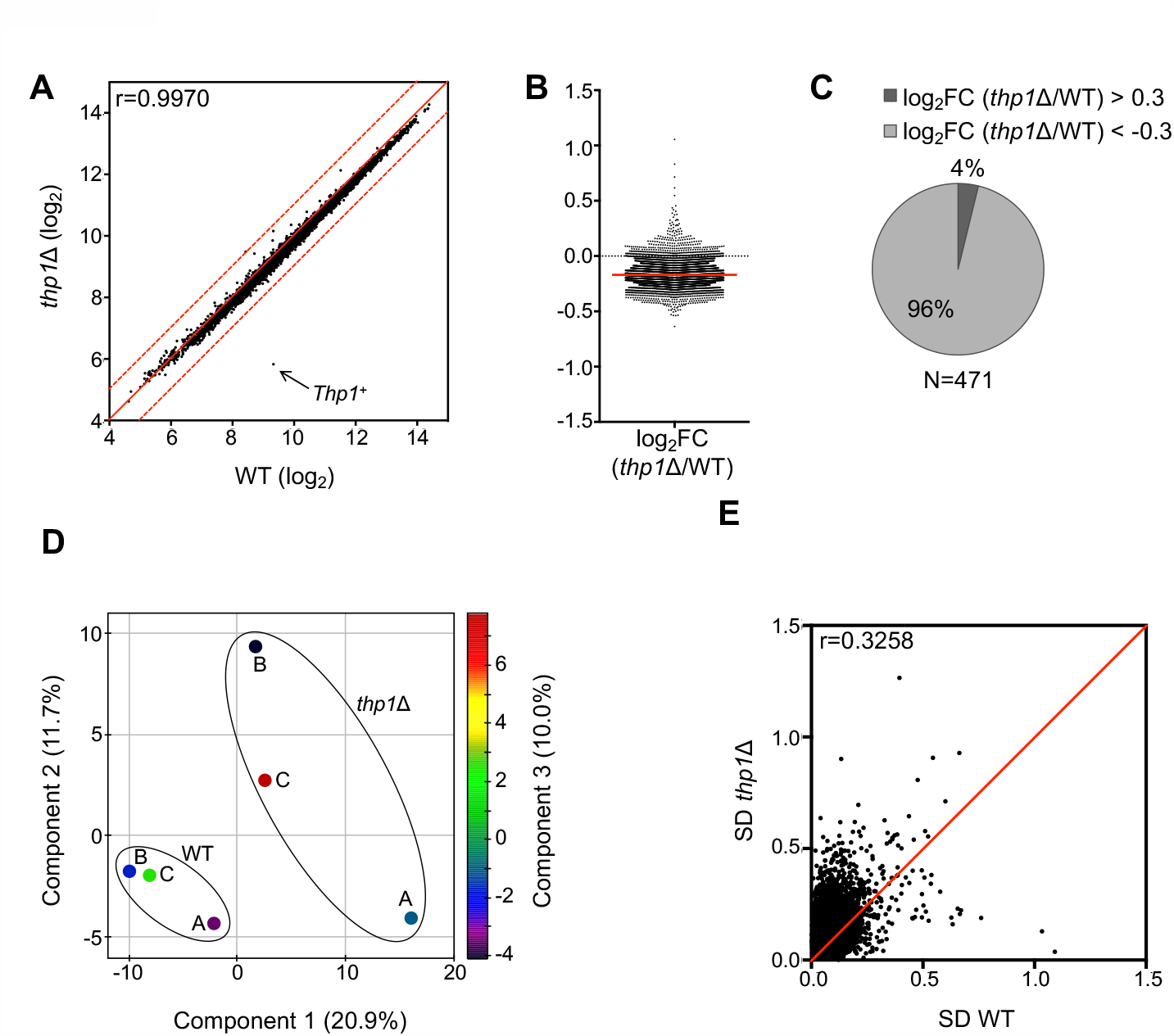
Gene expression in WT and Thp1 deficient cells. **(A)** Differential mRNA expression in WT and *thp1*Δ cells. Mean log_2_ expression values from wild-type triplicates was correlated with those of *thp1*Δ triplicates. The diagonal (red line), the Spearman correlation coefficient (r) and the point referring to the *tph1*^*+*^ (arrow) gene are indicated. **(B)** Relative gene expression in *thp1*Δ cells. mRNA expression values of *thp1*Δ were normalized to the wild-type (log_2_ fold change, log_2_FC), *thp1*^*+*^ itself was excluded from the analysis. The red line represents the median Log_2_FC. **(C)** A threshold of |log_2_| > 0.3 was applied to the normalized *thp1*Δ levels (N=471) and mRNAs were divided into up- and down-regulated genes. **(D)** Principal component analysis of single mRNA expression values (N=5022) of the wild-type and *thp1*Δ triplicates (A, B and C). **(E)** Comparison of standard deviations (SD) from wild-type and *thp1*Δ triplicates. SDs were calculated for each mRNA and *thp1*Δ SDs were plotted against WT SDs. The diagonal (red) and the Spearman correlation coefficient (r) are indicated.

A principal component analysis of mRNA expression patterns not only clearly separated the wild-type from *thp1*Δ samples but also revealed that the WT triplicate measurements were more closely related to each other than the *thp1*Δ triplicates (Fig. 5D). The higher variation among the *thp1*Δ triplicates was also visible in the standard deviations (SDs) resulting from the wild-type and *thp1*Δ expression measurements (Fig. 5E) and was generally present in all gene subsets analyzed Supplementary Fig. 2). Given that each of the replica was itself a pool of RNAs from three independent spore clones, this finding indicates a considerable degree of expression variability in cultures of Thp1 deficient cells. We tested whether the higher SD in the *thp1*Δ samples might be a technical consequence of the generally lower transcript signals obtained with these cells and found higher SDs in Thp1 deficient cell populations irrespective of whether the genes analyzed were down- or up regulated (Supplementary Fig. 3). Hence, the reduced and hyper-variable expression observed in Thp1-deficient cells appear to be a true biological effect.

## 4. Discussion

*S. pombe* has evolved with two active UDGs, Ung1 and Thp1, that seem to operate side by side. Their loss of function phenotypes show that, while both are capable of and even partially redundant in excising uracil from DNA, they do it in different ways. Uracil repair by Ung1 is efficient and productive, repair by Thp1 is inefficient and generates cytotoxic, mutagenic and recombinogenic intermediates. Hence, whereas Ung1 fulfills requirements of a robust DNA repair enzyme, Thp1 does not, and, therefore, likely has specialized functions beyond mutation avoidance, possibly in stabilizing gene expression.

In biochemical assays with cell-free extracts, Ung1 accounts for all detectable uracil excision activity in wild-type cells and the sensitivity of the assays used allows the conclusion that endogenous Thp1 accounts for less than 5% of total cellular uracil excision activity. Notwithstanding considerations of spatio-temporal separation of functions, this suggests that Ung1 provides the major activity for the repair of misincorporated dUMPs and deaminated cytosines. A similar prominent role has been reported on the grounds of biochemical evidence for mammalian UNG2 (58, 59). Contrary to this biochemical assessment, however, only the simultaneous inactivation of both, *thp1*^*+*^ and *ung1*^*+*^, gave rise to a significant accumulation of genomic uracil, showing that Thp1 does contribute to uracil repair and can compensate for the loss of Ung1 in living cells (Fig. 2B). These results suggest that Ung1 operates as the prime UDG in wild-type cells while Thp1 provides a backup activity, engaging mainly upon the loss or saturation of Ung1. Such redundancy also showed as a significant and synergistic increase in spontaneous mutation rates in *ung1*Δ *thp1*Δ double mutant cells, reflecting the loss of repair of deaminated cytosines.

We were surprised to find that *thp1*Δ *ung1*Δ double mutant cells show only a moderate (6.3-fold) increase in the C→T transition rate, this being assessed at an intragenic position of a transcribed gene. Provided that no other enzyme removes uracil from G•U mismatches in fission yeast, as implicated by the absence of additional UDG genes in the genome and the lack of residual uracil excision activity in double mutant cells, the C→T transition rate should equal the cytosine deamination frequency at this base pair. Hence, given a G-C content of 36% (60) and a genome size of 13.8 Mb, the genome-wide rate of endogenous cytosine deamination can be calculated to 0.04 events per haploid *S. pombe* cell cycle, i.e. one event per 25 cell divisions.

Notably, the canavanine forward mutation assay revealed a trend for increased mutations also in *thp1*Δ single mutant cells (Fig. 2D and 3D), implicating Thp1 in the repair of base lesions other than G•U mismatches. The spectra of the *ade6-M387* revertants in the *thp1*Δ background indicated that these lesions might generate C→G and C→A transversions. Etheno DNA adducts, in particular 3,N4-ethenocytosine, would represent a candidate lesion in this respect as it can give rise not only to C→T transitions but also to C→A transversions and was show to be a substrate for human TDG, *E. coli* Mug and Thp1 (36, 61, 62). Although the exact nature of the underlying Thp1-relevant base damage remains to be clarified, these small mutator effects suggest that proteins of the Mug family, including Thp1 and TDG, may indeed play a role in the defense against lipid-peroxidation mediated mutations.

Remarkably, exposing cells to the uracil analog 5-FU revealed diametrically opposite effects of Ung1- and Thp1-dependent repair. Whereas Ung1 was protective against 5-FU mediated cytotoxicity (Fig. 3B), Thp1 was detrimental, sensitizing wild-type as well as Ung1-deficient cells to the drug (Fig. 3C). A cell-sensitizing effect of Thp1 was also observed upon overexpression of AID. These effects demonstrate that uracil repair initiated by Thp1 is accompanied by the generation of cytotoxic repair intermediates. As previously noted for a 5-FU hyper-resistance observed upon *TDG* inactivation in mouse and human cells (24), base excision associated toxicity appears to be a consequence of particular enzymatic property of the Mug subfamily of UDGs, i.e. the rate limiting handover of AP-sites to downstream acting factors of BER (36, 63-65). Under conditions of uracil stress, such as under 5-FU treatment or AID expression, the engagement of Thp1 is thus likely to effect an accumulation of cytotoxic and mutagenic AP-sites, thereby compromising cell viability but also increasing mutation rates. Notably, 5-FU induced Thp1-dependent mutations were most pronounced in Ung1-deficient cells, showing that uracil repair by Ung1, in contrast to Thp1, is largely non-mutagenic.

AP-sites also interfere with replication fork progression (66-68), which can occasionally initiate homologous recombination. Our finding that 60% of spontaneous mitotic recombination at the *ade6* locus are dependent on Thp1 suggests that slow processing of spontaneously arising DNA base lesions represents a major source of gross genomic instability. In unchallenged cells, these can be AP-sites generated by Thp1 itself or arising from spontaneous base hydrolysis or the action of other DNA glycosylases. The latter is supported by the data showing that mitotic recombination induced by X-rays largely depends on Thp1. As the non-lethal dose applied in these experiments is expected to generate predominantly oxidative lesions, including substrates for other DNA glycosylases, this result suggests that Thp1 can interfere with the repair of any AP-site generated.

The mammalian Thp1 ortholog has specific functions in the control of gene activity at the level of DNA methylation and histone modifications (21, 25). We were not able to pinpoint a specific role of fission yeast Thp1 in gene regulation; Thp1-loss did not cause distinct patterns of transcriptional deregulation. Interestingly however, we found a widespread suppression of gene expression in Thp1-deficient cell populations. Also, gene expression was hyper-variable in the absence of Thp1, an observation that is rather robust, given that each replica analyzed contained pooled RNAs from three independent spore-derived cultures. These observations therefore indicate a contribution of Thp1 to the maintenance of stable gene activity. Given the absence of DNA methylation in fission yeast and, hence, a need for processing of demethylation intermediates (37, 69), Thp1 may operate through interactions with transcription factors or chromatin modifiers by a mechanism that remains to be determined. Interestingly, a TAP-tagging screening approach has identified a physical interaction of Thp1 with Cbf11 (70), a transcription factor capable of activating gene transcription (39).

While Ung1 fulfills expected requirements for an efficient DNA repair enzyme, Thp1 shows cytotoxic, mutagenic and recombinogenic activities that seem unsuitable for general genome maintenance. However, the functional properties of Thp1 may serve specific biological purposes under defined circumstances. One aspect to be considered is the temporal separation of function of the two UDGs in the cell cycle. Repair of uracil misincorporated during DNA replication would require an efficient, high fidelity pathway, whereas repair of uracils arising in non-replicating DNA would benefit from a more versatile UDG that facilitates coordination of base excision and AP-site release with the recruitment of downstream acting repair factors. Such functional separation was proposed for UNG2 and TDG in mammalian cells on the basis of differential and complementary expression of UNG2 and TDG during the cell cycle (14, 22). Addressing conditions other than vegetative growth, e.g. sporulation, pseudohyphal growth (71) or stationary phase, might thus reveal specific functions of Thp1. In addition, UDGs might act differentially in contexts where targeted genome and chromatin editing is required, such as somatic hypermutation, where targeted enzymatic deamination is coupled to error-prone BER (3, 72), or active DNA demethylation, where targeted DNA oxidation is linked to BER (9, 30). These are vertebrate examples and it is currently unclear whether and for what purpose analogous mechanisms operate in fission yeast. It is interesting, however, that the phenotype of the Thp1 knockout includes some perturbation of gene expression and it will be of interest to further identify potential Thp1-related gene regulatory mechanisms.

## ACKNOWLEDGEMENTS

We thank Marc Bühler, Jürg Kohli and Svend Petersen-Mahrt for valuable discussions, advice and the communication of unpublished data. We also thank Swend Petersen-Mahrt for the generous gift of AID antibodies.

