## Supplemental Information for "Uracil Repair - A Source of DNA Glycosylase Dependent Genome Instability"

### SUPPLEMENTARY METHODS

**Base release assay.** 60 bp double-stranded oligonucleotide substrates containing A•U and G•U mismatches were prepared by annealing an unlabeled oligonucleotide (5'-TAGACATTGCCCTCGAGGTACCATGGATCCGATGTC**(A/G)**ACCTCAAACCTAGACGAATTCCG-3') to a 5'-fluorescein (F)-labeled lower oligonucleotide strand (5'-F-CGGAATTCGTCTAGGTTTGAGGT**U**GACATCGGATCCATGGTACCTCGAGGGCAATGTCTA-3'). 0.5 µM labeled and 1 µM unlabeled oligonucleotides were annealed in 10 mM Tris-HCl pH 8.0 and 50 mM NaCl by heating to 95 °C for 5 min and gradual cooling to 25 °C over 30 min. Nicking assays were performed in a total volume of 20 µl nicking buffer (50 mM Tris-HCl pH 8.0, 1 mM DTT, 0.1 mg/ml BSA, 1 mM EDTA). 2 pmol of recombinant Thp1 or 7 µg of cell free extract was incubated with 2 pmol of substrate. When indicated, 1 unit of uracil-DNA glycosylase inhibitor (Ugi, New England BioLabs) was included. Reactions containing cell free extracts or recombinant Thp1 were incubated for 30 min at 30°C or for 15 min at 37°C, respectively. For AP-site cleavage, NaOH was added to a final concentration of 90 mM, followed by an incubation at 99°C for 10 min. DNA was ethanol-precipitated and boiled in 10 µl of formamide gel loading buffer (90% formamide, 1 X TBE) for 5 min at 99°C. After cooling, samples were separated using denaturing polyacrylamide gel electrophoresis. Fluorescein-labeled DNA was visualized using the blue fluorescent mode of the Storm 860 (Molecular Dynamics).

**DNA isolation in agarose plugs and PFGE**

DNA isolation in agarose plugs and PFGE were done as in {Baumann:2000uv} with some modifications (see Supplementary Methods). Cells were washed in water and resuspended in PRO buffer (1 M sorbitol, 25 mM EDTA, 20 mM Tris-HCl pH 8.0, 10 mM DTT) at a density of $5.5 \times 10^8$ cells/ml. $5 \times 10^8$ cells were treated with 1 mg/ml Zymolyase-20T (Amsbio) at 37°C for 60 min. Spheroplasts were collected and resuspended in 120 µl of TSE buffer (10 mM Tris-HCl pH 7.5, 900 mM sorbitol, 45 mM EDTA) before being mixed with 375 µl of 1.5% agarose (ultra pure L.M.P. agarose, GIBCO BRL) in TSE equilibrated at 43°C. Agarose plugs were poured, washed in PW1 (50 mM Tris- HCl pH 7.5, 250 mM EDTA, 1% SDS) at 50°C for 4 h and incubated twice in PW2 (10 mM Tris-HCl pH 9.0, 500 mM EDTA, 1% N-lauroyl sarcosine, 1 mg/ml proteinase K) at 50°C for 22 h. Finally, plugs were washed five times 60 min in 5 ml T10XE (10 mM Tris-HCl pH 7.5, 10 mM EDTA) and kept in T10XE at 4°C until use.

For digestion, plugs were washed twice in water for 15 min and once in 5 ml Ung-digestion buffer (20 mM Tris-HCl, pH 8.0, 150 mM NaCl) for 60 min at room temperature. Four quarters of the same plug were transferred to 4 ml Ung-digestion buffer supplemented with $MgCl_2$ (20 mM Tris-HCl, pH 8.0, 150 mM NaCl, 2 mM $MgCl_2$) containing either no enzyme(s), 13 units Ung uracil-DNA glycosylase (E. coli, New England BioLabs), 3.5 pmol APE1 (recombinant human AP-endonuclease) or both enzymes and incubated at 37°C for 4 h. The plugs were washed three times in 4 ml T10XE at 4°C, equilibrated two times for 1 h in 0.5 X TBE (pH 8.3, 45 mM Tris-HCL, 45 mM borate, 1 mM EDTA) and loaded onto 0.8% agarose gels (Chromosomal Grade Agarose, Bio-Rad) in 0.5 X TBE. Electrophoresis was performed in a CHEF DR III pulsed-field electrophoresis system (Bio-Rad) using 0.5 X TBE running buffer (72 h, 14°C, 2 V/cm, 1800 s switch time, included angle: 100°). DNA was stained with ethidium bromide (1 µg/ml) in 0.5 X TBE for 60 min and de-stained for 90 min in 0.5 X TBE.

### SUPPLEMENTARY FIGURES

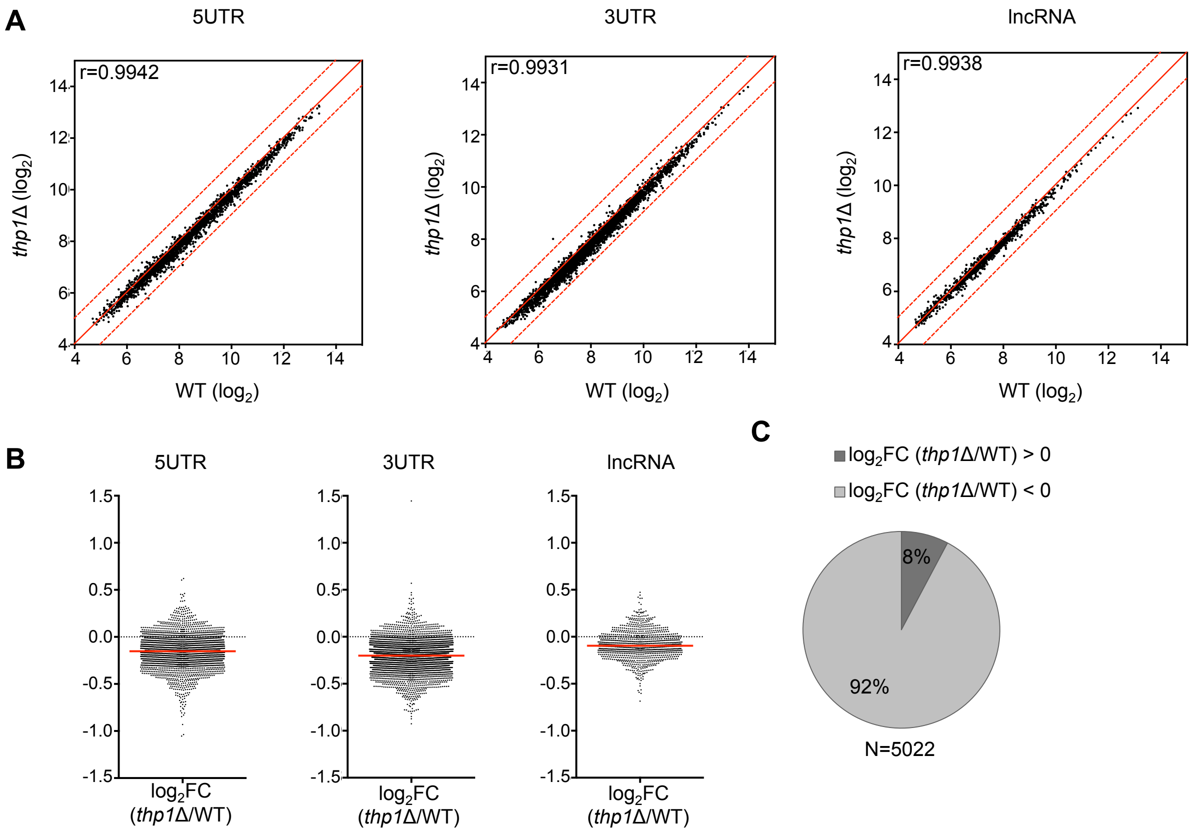


**Supplementary Figure 1 (related to Figure 5A-C). Expression of specific element classes in WT and *thp1Δ* cells.** 5′ UTR, 5UTR; 3′ UTR, 3UTR; long non-coding RNA, lncRNA. **(A)** Differential expression in WT versus *thp1Δ* cells. The diagonals (red) and the Spearman correlation coefficients (r) are indicated **(B)** Up- and down-regulation of gene expression in *thp1Δ* cells. 5UTR, 3UTR and lncRNA expression values of *thp1Δ* were normalized to the wild-type ($log_2$ fold change, $log_2FC$) and the value corresponding to the *thp1*$^+$ gene was omitted. Red line, median value. **(C)** The $Log_2FC$ (*thp1Δ*/WT) was calculated for all mRNAs that were sorted as up- and down-regulated.

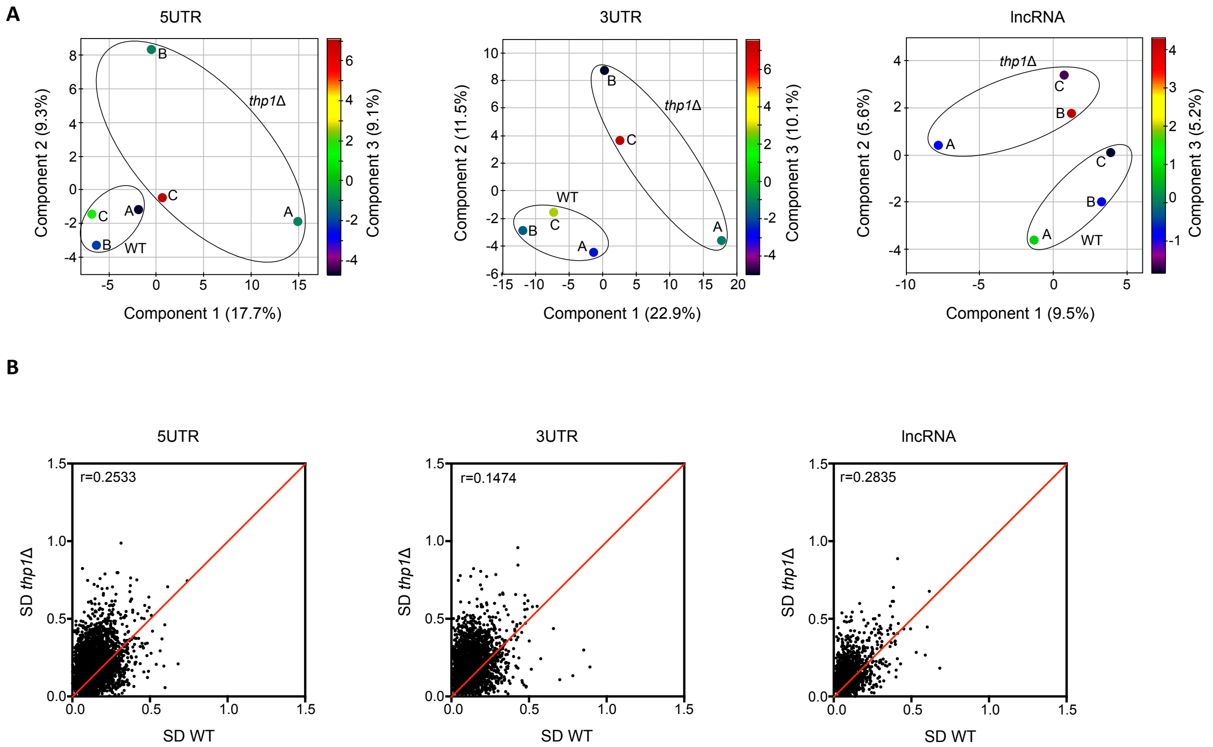


**Supplementary Figure 2 (related to Figure 5D to 5E). Variation between biological triplicates for 5UTRs, 3UTRs and lncRNAs. (A)** Principal component analysis of the six samples (3 wild-type and 3 *thp1*Δ samples). **(B)** Comparison of SDs from wild-type and *thp1*Δ triplicates. SDs were calculated for 5UTR, 3UTR or lncRNA elements and *thp1*Δ SDs were plotted against WT SDs. The diagonals (red) and the Spearman correlation coefficients (r) are indicated.

**A**

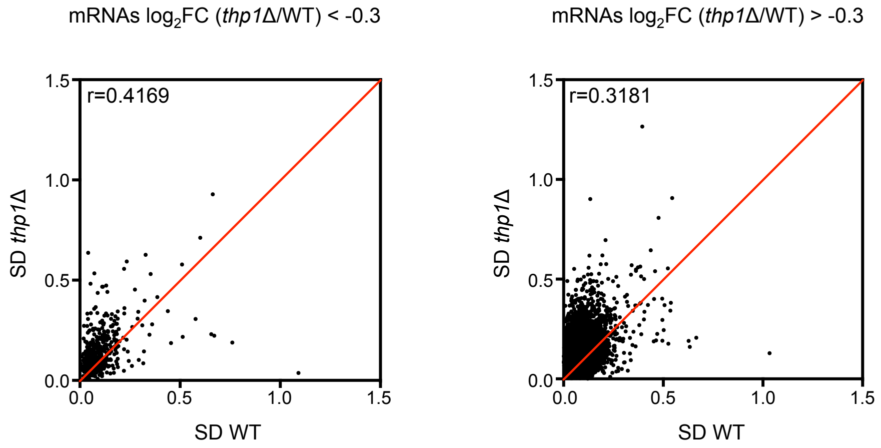


**B**

| | Average SD (WT) | Average SD (*thp1*Δ) | Average SD (WT/*thp1*Δ) |
|---|---|---|---|
| **log$_2$FC (*thp1*Δ/WT) <-0.3** | 0.153 | 0.219 | 1.426 |
| **log$_2$FC (*thp1*Δ/WT) >-0.3** | 0.113 | 0.174 | 1.537 |

**Supplementary Figure 3 (related to Figure 5E). SD comparison of mRNAs between WT and *thp1*Δ samples. (A)** All mRNAs were divided into relative enrichment above and below a log$_2$FC (*thp1*Δ/WT) of -0.3. WT SDs were correlated with *thp1*Δ SDs. The diagonals (red) and the Spearman correlation coefficients (r) are indicated. **(B)** Mean of SD for wild-type and *thp1*Δ as in (A).

**Supplementary Table 1.** Strains used in this study.

| Strain | Genotype | Figure | Source |
|---|---|---|---|
| 972 $h^-$ | wild-type *$h^-$* | 1,2,3 | (Leupold, 1954) |
| 975 $h^+$ | wild-type $h^+$ | 5, Supplementary 1-3 | (Leupold, 1954) |
| PRS 000g | *leu1-32 $h^+$* | 1,3,4 | lab stock |
| PRS 801 | *thp1::kanMX4 ung1::ura4 leu1-32 ura4-D18 $h^-$* | 1,3,4 | PRS000g x PRS571 |
| PRS 571 | *thp1::kanMX4 ung1::ura4 ura4-D18 $h^-$* | 1,2,3 | PRS556 x PRS603 |
| PRS 827 | *$thp1^+$-EGFP $h^-$* | 2 | PRS823 x PRS556 |
| PRS 555 | *thp1::kanMX4 $h^-$* | 2,3,5, Supplementary 1-3 | 975$h^+$ x PRS550 |
| PRS 603 | *ung1::ura4 ura4-D18 $h^-$* | 2,3 | this study |
| PRS 053 | *ade6-M387 ura4-D18 $h^-$* | 2 | lab stock |
| PRS 563 | *thp1::kanMX4 ade6-M387 ura4-D18 $h^-$* | 2 | this study |
| PRS 605 | *ung1::ura4 ade6-M387 ura4-D18 $h^-$* | 2 | this study |
| PRS 575 | *thp1::kanMX4 ung1::ura4 ade6-M387 ura4-D18 $h^-$* | 2 | this study |
| PRS 807 | *ura4-D18 ade6-M375 $h^-$ int::pUC8/ura4/ade6-L469* | 4 | this study |
| PRS 809 | *thp1::kanMX4 ura4-D18 ade6-M375 $h^-$ int::pUC8/ura4/ade6-L469* | 4 | this study |
| PRS 811 | *ung1::ura4 ura4-D18 ade6-M375 $h^-$ int::pUC8/ura4/ade6-L469* | 4 | this study |
| PRS 813 | *thp1::kanMX4 ung1::ura4 ura4-D18 ade6-M375 $h^-$ int::pUC8/ura4/ade6-L469* | 4 | this study |
| PRS 823 | *ura4-D18 $thp1^+$::3'-$egfp^+$ $h^-$* | 2 | this study |
| LV10 | *rec12::ura4 ura4-D18 leu2-112 ade6-M26 $h^+$* | 3 | this study |
| LV22 | *thp1::kanMX4 rec12::ura4 ura4-D18 leu2-112 ade6-M26 $h^-$* | 3 | this study |
| LV40 | *thp1::kanMX4 ung1::ura4 rec12::ura4 ura4-D18 leu2-112 ade6-469 $h^+$* | 3 | this study |
| LV41 | *ung1::ura4 rec12::ura4 ura4-D18 leu2-112 ade6-M26 $h^+$* | 3 | this study |
